# Drug Interactions Between Dolutegravir and Artemether-Lumefantrine or Artesunate-Amodiaquine

**DOI:** 10.1101/351684

**Authors:** Stephen I Walimbwa, Mohammed Lamorde, Catriona Waitt, Julian Kaboggoza, Laura Else, Pauline Byakika-Kibwika, Alieu Amara, Joshua Gini, Markus Winterberg, Justin Chiong, Joel Tarning, Saye H Khoo

## Abstract

Across sub-Saharan Africa, patients with HIV on antiretrovirals often get malaria and need cotreatment with artemisinin-containing therapies. We undertook two pharmacokinetic studies in healthy volunteers, using standard adult doses of artmether-lumefantrine (AL) or artesunate-amodiaquine (AS-AQ) given with 50mg once daily dolutegravir (DTG) to investigate the drug-drug interaction between artmether-lumefantrine or artesunate-amodiaquine and DTG. The DTG/artmether-lumefantrine interaction was evaluated in a two-way cross-over study and measured artemether (ARM), dihydroartemisinin (DHA), lumefantrine (LF), desbutyl-lumefantrine (DBL) over 264h. The DTG/artesunate-amodiaquine interaction was investigated using a parallel study design due to long half-life of the amodiaquine metabolite, desethylamodiaquine (DEAQ) and measured artesunate (ARS), amodiaquine (AQ), DEAQ over 624h. Non-compartmental analysis was performed, and geometric mean ratios and 90% confidence intervals generated for evaluation of both interactions. Dolutegravir did not significantly change the maximum concentration in plasma, time to maximum concentration and area under the concentration-time curve (AUC) for ARM, DHA, LF and DBL nor significantly alter AUC for ARS, DHA, AQ and DEAQ. Co-administration of dolutegravir with AL resulted in a 37% decrease in DTG trough concentrations. Co-administration of dolutegravir with AS-AQ resulted in a decrease of approximately 42% and 24% in DTG trough concentrations and AUC respectively. Study drugs were well-tolerated with no serious adverse events. Standard doses of artmether-lumefantrine and artesunate-amodiaquine should be used in patients receiving DTG. The significant decreases in DTG trough concentrations with artemether-lumefantrine and artesunate-amodiaquine and DTG exposure with artesunate-amodiaquine are unlikely to be of clinical significance as DTG trough concentrations were above DTG target concentrations of 64ng/mL.

## INTRODUCTION

Over 90% of malaria cases occur in sub-Saharan Africa (SSA) the region with the greatest burden of HIV(1). Drug-drug interactions (DDIs) between antiretrovirals and artemisinin-based combination therapies (ACT) frequently occur and may affect the clinical effectiveness of commonly utilized antimalarials artemether-lumefantrine (AL) and artesunate-amodiaquine (AS-AQ)(2–4).The likely adoption of dolutegravir (DTG) in preferred first-line antiretroviral therapy (ART) regimens(5) makes a DDI study with antimalarials an urgent priority. Dolutegravir is predominately metabolized via uridine diphosphate glucuronyl transferase 1A1 (UGT 1A1) with minor input from cytochrome P450 (CYP)-3A4(6).

Both artemether (ARM) and lumefantrine (LF) are predominantly metabolized via CYP3A4, CYP2B6, CYP2C9 and CYP2C19 to active metabolites dihydroartemisinin (DHA) and desbutyl-lumefantrine (DBL) respectively. Artesunate (AS) is a prodrug and substrate of CYP2A6 and undergoes rapid hydrolysis to DHA while amodiaquine (AQ) is extensively metabolized by CYP 2C8 to its active metabolite, N-desethylamodiaquine (DEAQ)(5,7). Co-administration of artemether-lumefantrine with inducers of CYP3A4 results in significant reductions in artemether and dihydroartemisinin exposures(8). Similarly, clinically significant DDIs with ritonavir-boosted protease inhibitor ART regimens have been reported(3). However, data for dolutegravir are lacking. We investigated the pharmacokinetic (PK) interactions between dolutegravir with artemether-lumefantrine or artesunate-amodiaquine and assessed safety and tolerability of the drug combinations.

## METHODS

### Ethics

The study was approved by the Joint Clinical Research Centre Institutional Review Board, Kampala, Uganda and University of Liverpool Research Ethics Committee, Liverpool, UK and registered on ClinicalTrials.gov (NCT 02242799). The study was conducted in compliance with International Council for Harmonisation Good Clinical Practice guidelines, the current ethical principles in the Declaration of Helsinki and applicable local regulatory requirements.

### Study Design

Two open-label, fixed sequence studies between DTG and artemether-lumefantrine (Study A), or artesunate-amodiaquine (Study B) were conducted at the Infectious Diseases Institute, Kampala, Uganda. Inclusion of 16 subjects in Study A was calculated to have a >80% power to detect a change in area under the concentration-time curve (AUC) outside FDA limits for bioequivalence for dolutegravir and lumefantrine (assuming coefficient of variation ≤30%), and to detect a ≥32% change in dihydroartemisinin levels. Including 30 subjects in Study B would yield an 80% power to detect an AUC difference of >25-30% (DTG and DEAQ), and a ≥42% change for dihydroartemisinin.

### Eligibility Criteria

Consenting healthy adults (≥18 years) weighing above 40 kg, without malaria and HIV were eligible to participate if they had no evidence of systemic disease, were willing to use mosquito bed nets and able to comply with study procedures. Pregnant or breastfeeding women and female volunteers unwilling to use reliable contraception during the study were also excluded.

### Dosing and Sampling

#### Study A (Artemether-lumefantrine)

Study A used a random sequence two-way crossover study design with participants randomized to Arm 1 or Arm 2. In Arm 1, participants received six doses of oral artemether-lumefantrine (80/480 mg) tablets over three days (regimen used for treatment of uncomplicated malaria) with PK sampling at 0,1,2,3,4,8,12,24,48,72,96,168 and 264 hours after the final dose. After a 21 day washout, they received dolutegravir alone for six days with PK sampling on day 6 at 0, 1,2,3,4,8,12 and 24 hours post-dosing. Subsequently they received three days of twice daily AL plus dolutegravir, with PK sampling at 0,1,2,3,4,8,12,24,48,72,96,168 and 264 hours after the final doses of both drugs using the sampling time points described. In Arm 2, participants received the dolutegravir and DTG-AL combination with PK sampling as detailed for Arm 1, followed by artemether-lumefantrine alone after the 21 day washout period.

#### Study B (Artesunate-amodiaquine)

Study B used a parallel group design due to the long half-life of the amodiaquine active metabolite DEAQ. Participants were randomized to receive AS-AQ (4 mg/Kg AS, 10 mg/kg AQ) once daily for three days with PK sampling at 0,1,2,3,4,8,12,24,48,72,96,120,228 and 624 hours post-last dose (Arm 1), or dolutegravir for seven days with PK sampling at 0,1,2,3,4,8,12 and 24 hours after the last dose, followed by AS-AQ once daily together with dolutegravir once daily for three days, with PK sampling after the last dose of both drugs using the sampling time points listed (Arm 2).

### Safety assessments

At screening, a medical history, physical examination, urine pregnancy test, rapid malaria and HIV tests and safety bloods (hemoglobin, white cell count, platelets, urea, creatinine, electrolytes, ALT) were performed. A 12-lead ECG was performed at screening, intensive PK and at end of study. Safety bloods were repeated at every intensive PK visit and prior to discharge from the study. Laboratory and clinical abnormalities were graded for severity according to the U.S National Institutes for Health Division of AIDS (DAIDS) Table for Grading Severity of Adult and Pediatric Adverse Events.

### Pharmacokinetic analysis

Dolutegravir blood samples were collected in ethylenediaminetetraacetic acid (EDTA) tubes(9). Samples for artemisinin-based combination therapies were collected in either lithium heparin or fluoride-oxalate tubes to minimize ex-vivo degradation of artemisinins to DHA by plasma esterases(10). Blood samples were delivered within 15 minutes of collection to the laboratory for separation and storage at −80°C until shipment to the Liverpool Bioanalytical Facility and Mahidol University for quantification of dolutegravir and ACTs, respectively. Both laboratories participate in external Quality Assurance programmes and operate to GCP with assays validated according to published FDA guidelines.

Dolutegravir was extracted using liquid-liquid extraction and analyzed using a validated reversed phase liquid chromatography with a lower limit of quantification (LLOQ) set at 10 ng/ml and precision of 5% at low QC (30ng/mL)(9).

Antimalarial drugs were extracted using solid-phase extraction, and quantified by liquid chromatography tandem-mass spectrometry (LC-MS/MS). For artemether/dihydroartemisinin the total assay coefficient of variation was < 6% with LLOQ of 1.14 ng/ml. For artesunate/ dihydroartemisinin the coefficient of variation was < 7% with LLOQ of 0.119 ng/ml (AS) and 0. 196 ng/ml (DHA)(10). For lumefantrine/desbutyl-lumefantrine total coefficient variation was <6% with LLOQ of 7.77 ng/ml (LF) and 0.81 ng/ml (DBL)(11). For amodiaquine/N-desethylamodiaquine the total coefficient of variation was <8% with LLOQ of 0.86 ng/ml (AQ) and 1.13 ng/ml (DEAQ).

### Statistical analysis

Pharmacokinetic parameters including the area under the concentration-time curve to the last measurable time point (AUC_0-t_), terminal elimination half-life (t_1/2_), maximum concentration (C_max_) and time to C_max_ (T_max_) were estimated using non-compartmental analysis (WinNonlin, Phoenix,version 6.1, Pharsight, Mountain View, CA). PK data were log-transformed to calculate geometric mean ratios (GMR), with 90% CI evaluated using paired (Study A) or unpaired (Study B) t-tests and back-transformed to absolute ng/mL concentrations. An analysis of variance (ANOVA) was performed by SPSS (Windows Standard version 22; SPSS, Inc., Chicago, IL) on PK parameters (AUC_0-t_, C_max_, C_24_) using generalized linear models procedures to assess potential sequence and period related effects.

## RESULTS

Forty-eight participants were enrolled into both studies of whom 39 completed study procedures. Demographic variables of participants who completed study procedures are presented in Table 1.

**Table 1.** Participant median (IQR) baseline demographic variables

| Parameter (n=39) | Study A (n=14) |  | Study B (n=25) |  |
| --- | --- | --- | --- | --- |
|  | Sequence1(n=7) | Sequence 2 (n=7) | Arm 1 (n=13) | Arm 2 (n=12) |
| Age (years) | 29 (21-32) | 25 (23-29) | 24 (23-28) | 30.5 (23.5-34) |
| Weight (Kg) | 55.5 (54-64) | 59 (54-62) | 59.5 (57-65) | 60.25 (58-68.25) |
| BMI (Kg/m<sup>2</sup>) | 21.1 (17-22.8) | 21.2 (20.1-21.5) | 21.4 (19.8-24.5) | 20.5 (18.95-24.5) |
| Haemoglobin (g/dL) | 15.1 (13.3-17.7) | 14.7 (13.7-17.0) | 15.2 (13.5-16.2) | 15.3 (14.5-16.3) |
| ALT (IU/L) | 12 (9-20) | 15 (12-17) | 15 (13-19) | 18 (13-20) |
| Total bilirubin (mg/dL) | 0.7 (0.4-0.9) | 0.6 (0.4-1.1) | 0.5 (0.3-0.7) | 0.6 (0.35-1.65) |
| Urea (mg/dL) | 7 (6-9) | 7 (6-9) | 7 (5-8) | 8 (6.5-11) |
| Creatinine (mg/dL) | 0.76 (0.59-0.93) | 0.75 (0.62-0.79) | 0.82 (0.67-0.92) | 0.85 (0.69-0.92) |
| Corrected QT*interval (ms) | 387 (378-407) | 415 (397-429) | 396 (369-408) | 400.5 (373.5-414.5) |

*Note: QTc\*- Fridericia's formula*
*Abbreviations: IQR, interquartile range; BMI, body mass index; ALT, alanine transaminase*

### Antimalarial Pharmacokinetics

#### Effect of dolutegravir on artemether-lumefantrine pharmacokinetics (Study A)

In study A, 14 participants received AL (7 Arm 1; 7 Arm 2). The artemether/dihydroartemisinin PK profiles (0-24 hours), lumefantrine/desbutyl-lumefantrine PK profiles (0-264 hours), and associated PK parameters are presented in Table 2.

**Table 2.**
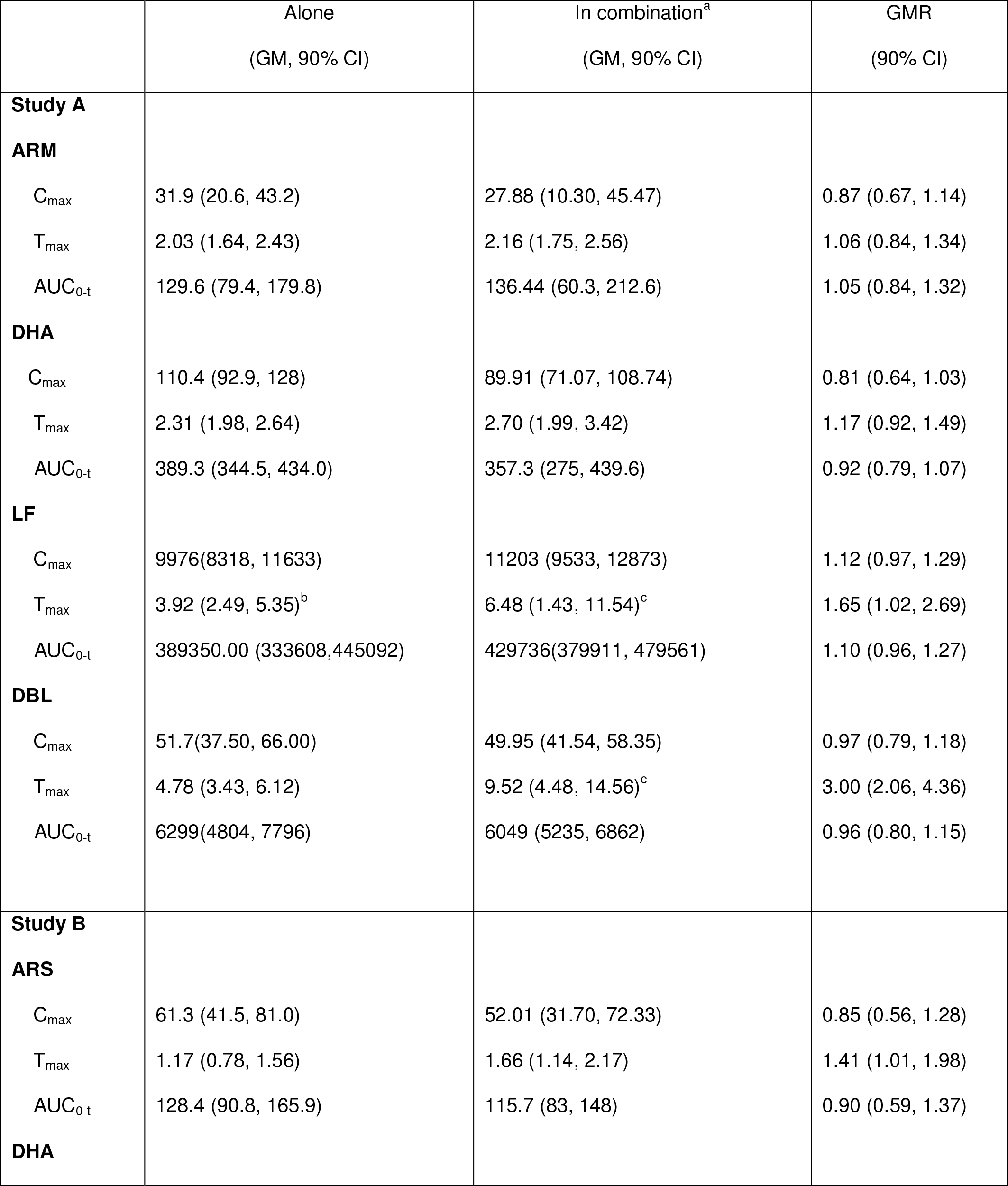
Artemether-lumefantrine, artesunate-amodiaquine pharmacokinetic parameters alone and in combination with dolutegravir

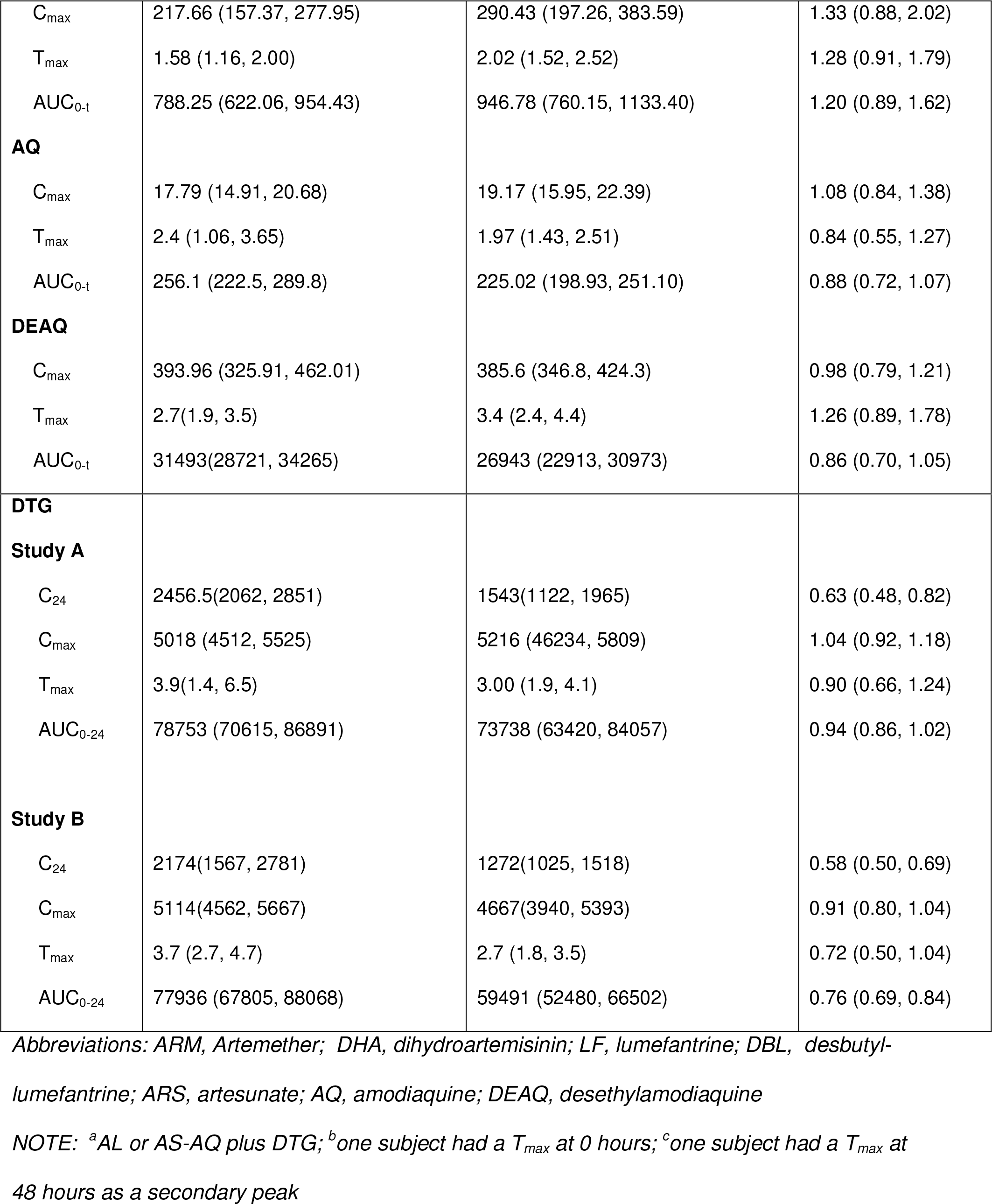

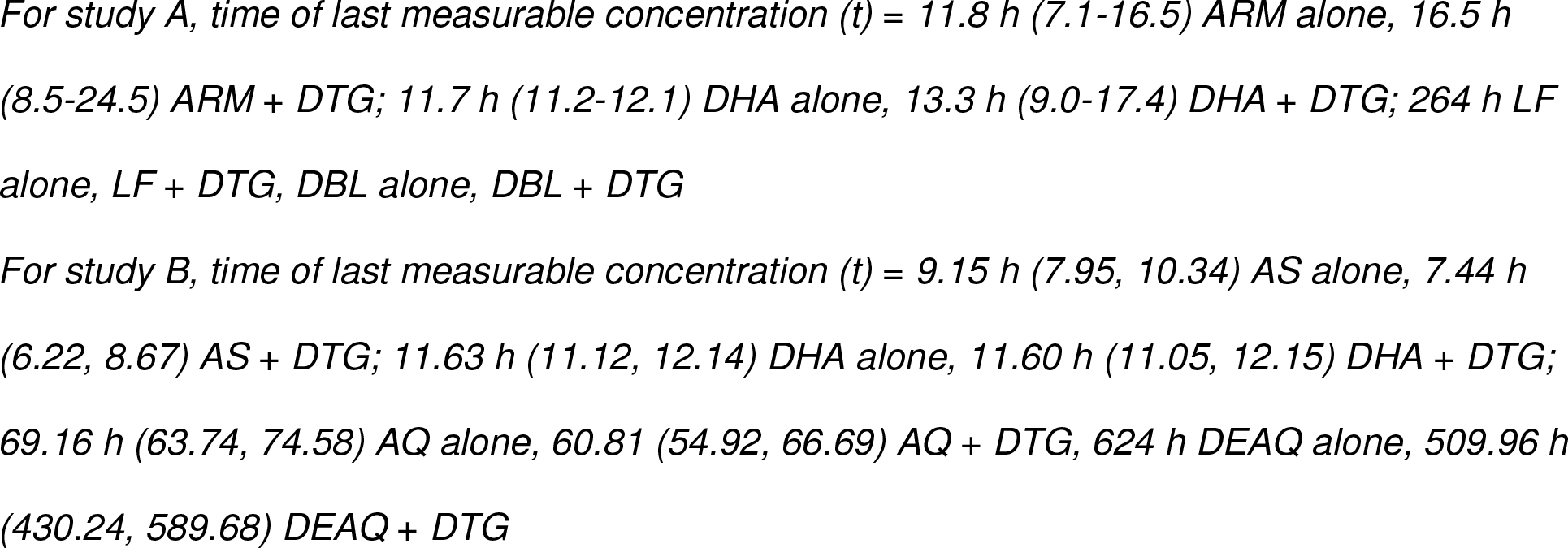

When artemether-lumefantrine was administered alone, geometric mean (GM 90% CI) artemether maximum concentrations of 31.9 ng/ml (20.6-43.2) were reached on average in 2 hours, with AUC_0-t_ of 129.6 ng.h/ml (79.4-179.8). The active metabolite DHA achieved peak concentrations of 110.4 ng/ml (92.9-128.0) after 2.3 hours with AUC_0-t_ of 389.3 ng.h/ml (344.5-434.0). Artemether and dihydroartemisinin were eliminated from plasma with an average halflife of 5 and 2.5 hours, respectively.

Lumefantrine showed peak concentrations approximately four hours after drug administration, with a C_max_ and AUC_0-t_ of 9976 ng/ml (8318-11633) and 389350 ng.h/ml (333608-445092), respectively. The lumefantrine metabolite DBL had a C_max_ and AUC_0-t_ of 51.7 ng/ml (37.5-66.0) and 6699 ng.h/ml (4804-7796) respectively, representing approximately 1.7% of total circulating lumefantrine. The elimination half-life of lumefantrine and desbutyl-lumefantrine were 83 and 142 hours, respectively.

The GMR for each antimalarial and metabolite are presented in Table 2. Co-administration of artemether-lumefantrine with DTG did not significantly alter C_max_, AUC_0-t_ or clearance of artemether, lumefantrine or their metabolites. Furthermore, the time of the last measurable concentration (t) for all artemether-lumefantrine components did not significantly differ when administered alone or in combination with DTG. The analysis of variance (ANOVA) showed no evidence of a significant sequence or period effect upon artemether-lumefantrine PK.

### Effect of dolutegravir on artesunate-amodiaquine pharmacokinetics (Study B)

In study B, 25 participants received AS-AQ. 13 subjects in Arm 1 were administered artesunate-amodiaquine alone, and 12 subjects in Arm 2 were administered artesunate-amodiaquine with DTG. The artesunate/ dihydroartemisinin PK profiles (0-12 hours), amodiaquine/N-desethylamodiaquine PK profiles (0-624 hours), and associated PK parameters are presented in Table 2.

When artesunate-amodiaquine was administered alone, maximum artesunate concentrations of 61.3 ng/ml (41.5-81.0) were reached within 1.2 hours, with an overall AUC_0-t_ of 128.4 ng.h/ml (90.8-165.9). Dihydroartemisinin exposures were on average 6-fold higher than corresponding artesunate AUC_0-t_ values. Artesunate and dihydroartemisinin geometric mean half-life was 1.9 and 2.2 hours, respectively. Similarly, amodiaquine was rapidly absorbed (T_max_ = 2.4 hours) and was detectable in plasma for approximately 70 hours post-dose; overall amodiaquine AUC_0-t_ was 256.1 ng.h/ml (222.5-289.8). Amodiaquine was rapidly and extensively converted to DEAQ (T_max_ = 2.7 hours); N-desethylamodiaquine exposures were approximately 122-fold higher than amodiaquine, and persisted in plasma for up to 624 hours post-dose (terminal elimination halflife ~10 days).

Co-administration of DTG with artesunate-amodiaquine did not significantly alter the AUC_0-t_ for AS (p=0.77), DHA (p=0.27), AQ (p=0.14) or DEAQ (p=0.69).

### Dolutegravir pharmacokinetics

Dolutegravir was administered with and without antimalarials to 14 participants in study A (7 Arm 1; 7 Arm 2) and 12 participants in study B (Arm 2). Individual PK profiles, geometric mean (90% CI) dolutegravir PK profiles (0-24 hours) for study A and B are depicted in Figure 1 and Figure 2 respectively.

**Figure 1.**
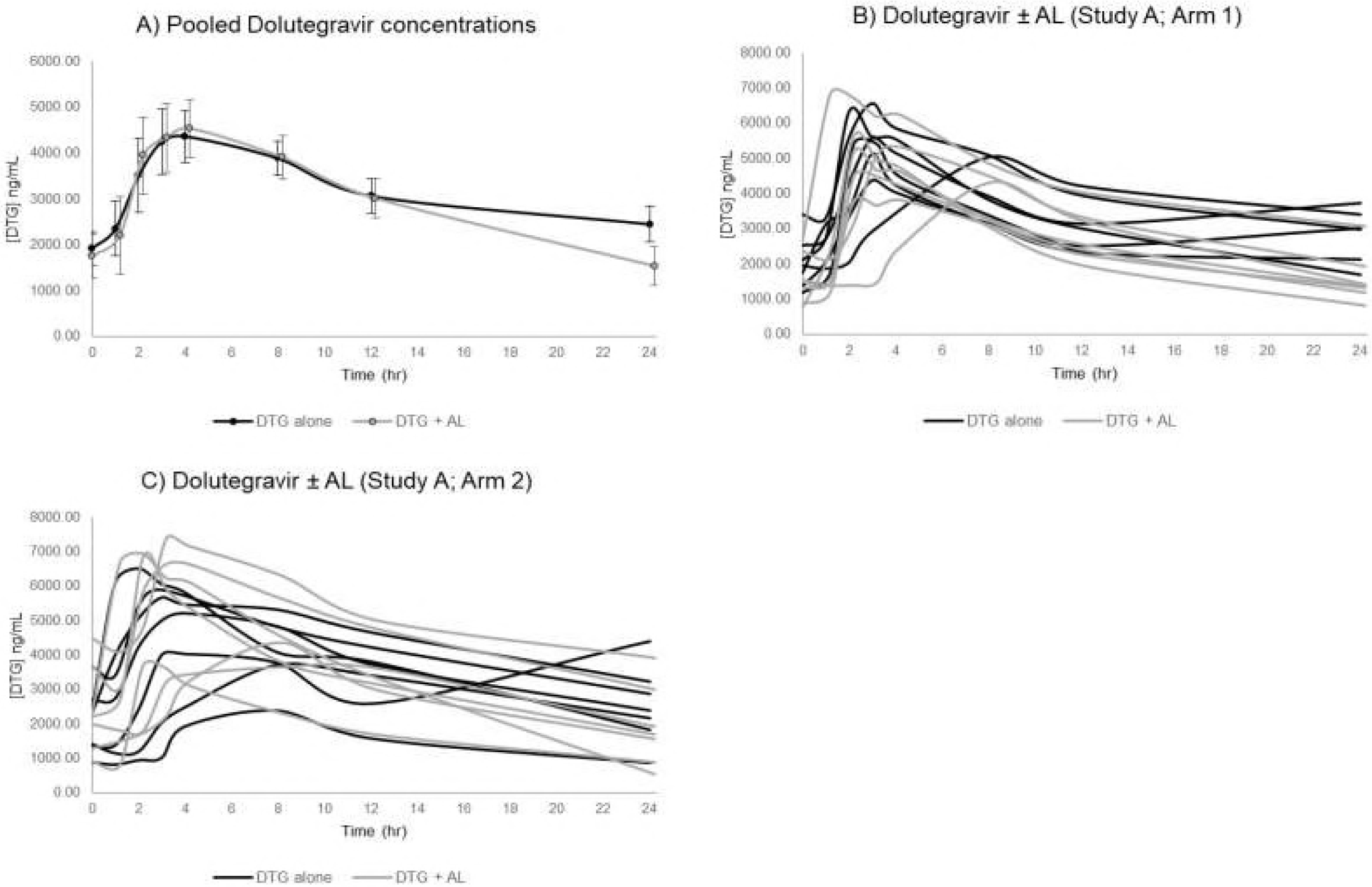
Study A (DTG+AL) pharmacokinetics

**Figure 2.**
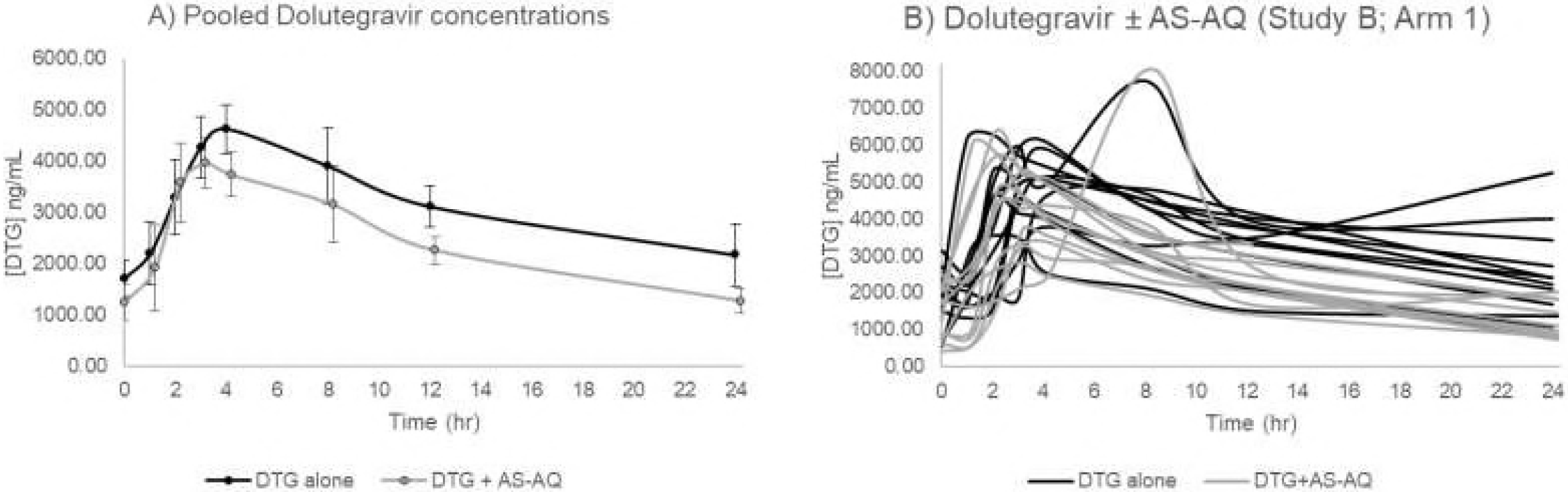
Study B (DTG ± AS-AQ) pharmacokinetics

### Effect of artemether-lumefantrine on dolutegravir pharmacokinetics (Study A)

In study A, dolutegravir C_max_ of 5018 ng/ml (4512-5525) was reached at 3.9 hours post-dose, with an overall AUC_0-24_ of 78753.4 ng.h/ml (70615-86891).

Co-administration of DTG with artemether-lumefantrine resulted in a 37% decrease in dolutegravir C_24_ [GMR = 0.63 (0.48-0.82)]. No significant changes were observed in dolutegravir AUC_0-24_ or C_max_ when DTG was administered with artemether-lumefantrine (Table 2). The ANOVA revealed no significant sequence effect upon dolutegravir PK. However, there was a significant period effect (DTG alone vs. DTG + AL) for dolutegravir C_24_ in both arms (p=0.025).

### Effect of artesunate-amodiaquine on dolutegravir pharmacokinetics (Study B)

In study B, dolutegravir C_max_ of 5114 ng/ml (4562-5667) was reached at 3.7 hours post-dose, with an overall AUC_0-24_ of 77936 ng.h/ml (67805-88068).

Co-administration of DTG with artesunate-amodiaquine resulted in a significant decrease of approximately 42% and 24% in dolutegravir C_24_ and AUC_0-24_, respectively as presented in Table 2.

### Safety

All adverse events were Grade 1 or 2 in severity. Gastro-intestinal adverse events were more frequent among participants receiving artesunate-amodiaquine.

## DISCUSSION

This is the first study to examine for drug interactions between dolutegravir with artemether-lumefantrine and artesunate-amodiaquine. We found no significant impact of dolutegravir upon drug exposure of either antimalarial regimen, suggesting that standard doses of artemether-lumefantrine and artesunate-amodiaquine should be used when co-administered with dolutegravir. These findings are important given the increasing use of dolutegravir in first-line antiretroviral therapy regimens. The results confirm the low propensity for dolutegravir to perpetrate clinically significant DDIs, as judged by in-vitro observations of minimal effects on drug transporters and cytochrome P450 enzymes(6,12).

We observed that artemether-lumefantrine was not associated with any significant change in dolutegravir exposure parameters (C_max_, AUC_0-24_). However, dolutegravir C_24_ was 37% lower with artemether-lumefantrine than when given alone. The reasons for this are unclear but may have been driven in part by an unexplained rise in dolutegravir C_24_ in some participants or weak induction of CYP3A4 by artemether and dihydroartemisinin. Additional intake of dolutegravir prior to the last sampling point at 24 hours was unlikely, as subjects were instructed not to take the next dose before this time-point, and were issued with an exact number of pills which precluded such additional intake.

With artesunate-amodiaquine, we observed an unexplained statistically significant reduction of 42% and 24% for dolutegravir C_24_ and AUC_0-24_ respectively. However, in all subjects who received DTG with artemether-lumefantrine or artesunate-amodiaquine, the dolutegravir C_trough_ was comparable to or above 1100ng/mL, the mean C_trough_ observed in prior dolutegravir phase 3 adults trials. The target minimum effective concentrations for dolutegravir are unknown, although a DTG protein-adjusted IC_90_ of 64ng/mL has been proposed. In a phase II study, C_trough_ concentrations over 324ng/mL after 10 days of DTG monotherapy were associated with virological efficacy(12). All subjects in our study had C_trough_ concentrations exceeding these targets, suggesting that the modest pharmacokinetic changes observed have unlikely clinical significance, especially given the short duration of antimalarial therapy.

Surprisingly, dolutegravir concentrations in this study of black African healthy volunteers were somewhat higher than previously reported in Caucasians. It should be noted that dolutegravir was dosed with a moderate fat meal, and concentrations observed are consistent with reports on the food effect upon DTG bioavailability(13).

The combination of dolutegravir with artemether-lumefantrine and artesunate-amodiaquine was well tolerated. Nausea, the most common study drug-related adverse event, was reported predominately in the artesunate-amodiaquine arm. This safety profile was consistent with the approved drug labelling (13–15).

In conclusion, standard treatment regimens of artesunate-amodiaquine, and of artemether-lumefantrine should be prescribed when treating malaria in HIV-infected patients receiving a dolutegravir based antiretroviral regimen.

## ACKNOWLEDGMENTS

The authors thank the study participants and members of the DolACT clinical trial team (R. Nakijoba, F. Aber, J. Magoola, E. Ssempijja). We also thank the staff of the Infectious Diseases Institute research department (H. Onen, R. Nalumenya, S. Nabukenya). We acknowledge the contribution of the trial steering committee and IDSMB (Ed Wilkins, David Burger, and Victor Mwapasa)

## Meeting presentation

Conference on Retroviruses and Opportunistic Infections; March 04 to 7, 2018; Boston; MA (Abstract # 459)

## Transparency declarations

SK has received research funding from Merck, Gilead and ViiV Healthcare. The Liverpool HIV Drug Interactions website (www.hiv-druginteractions.org) has received support from ViiV, Gilead Sciences, Merck, and Janssen. All other authors: none to declare.

---

**FIG S1.**
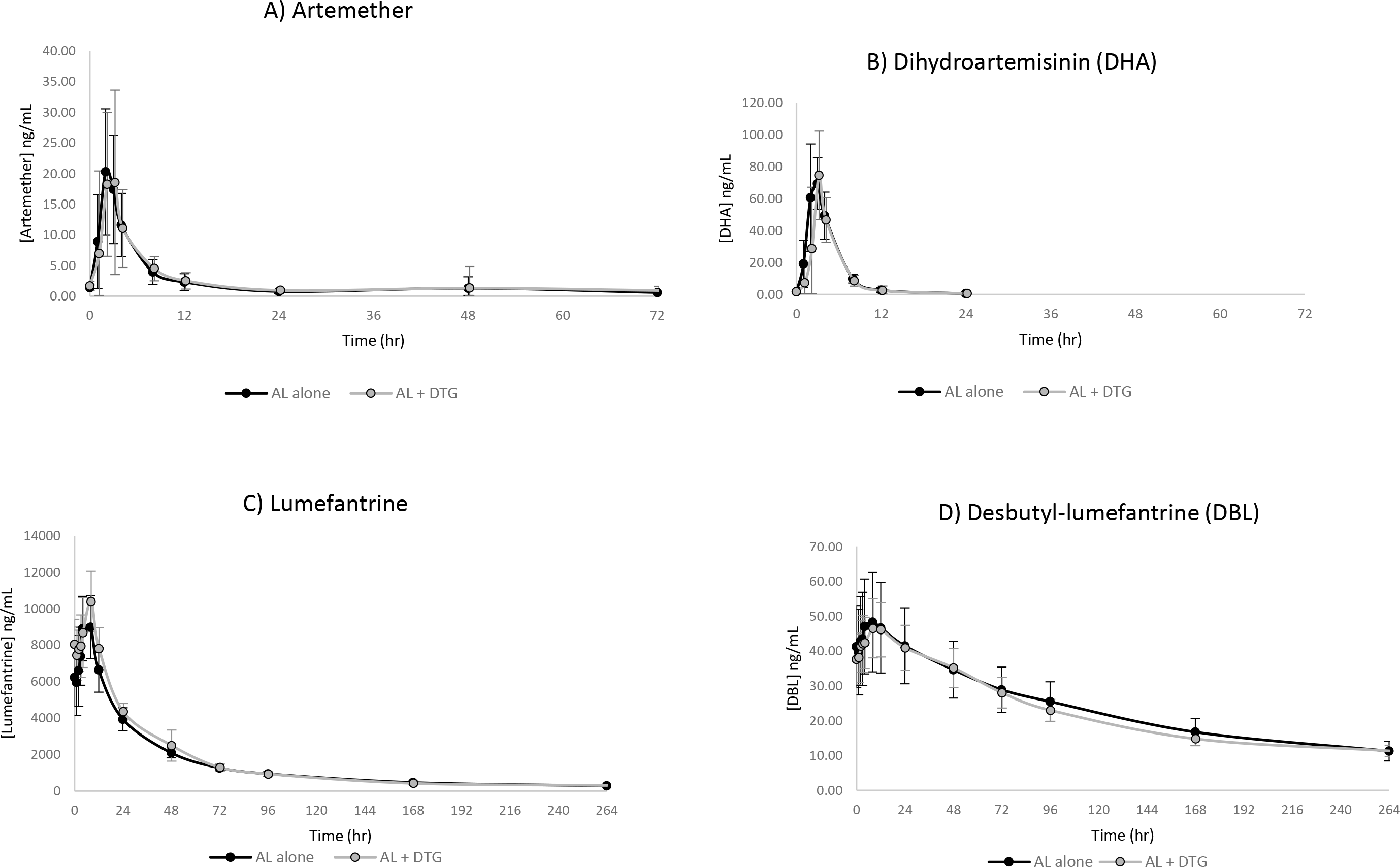
Artemether-lumefantrine parent and active metabolite PK.

**FIG S2.**
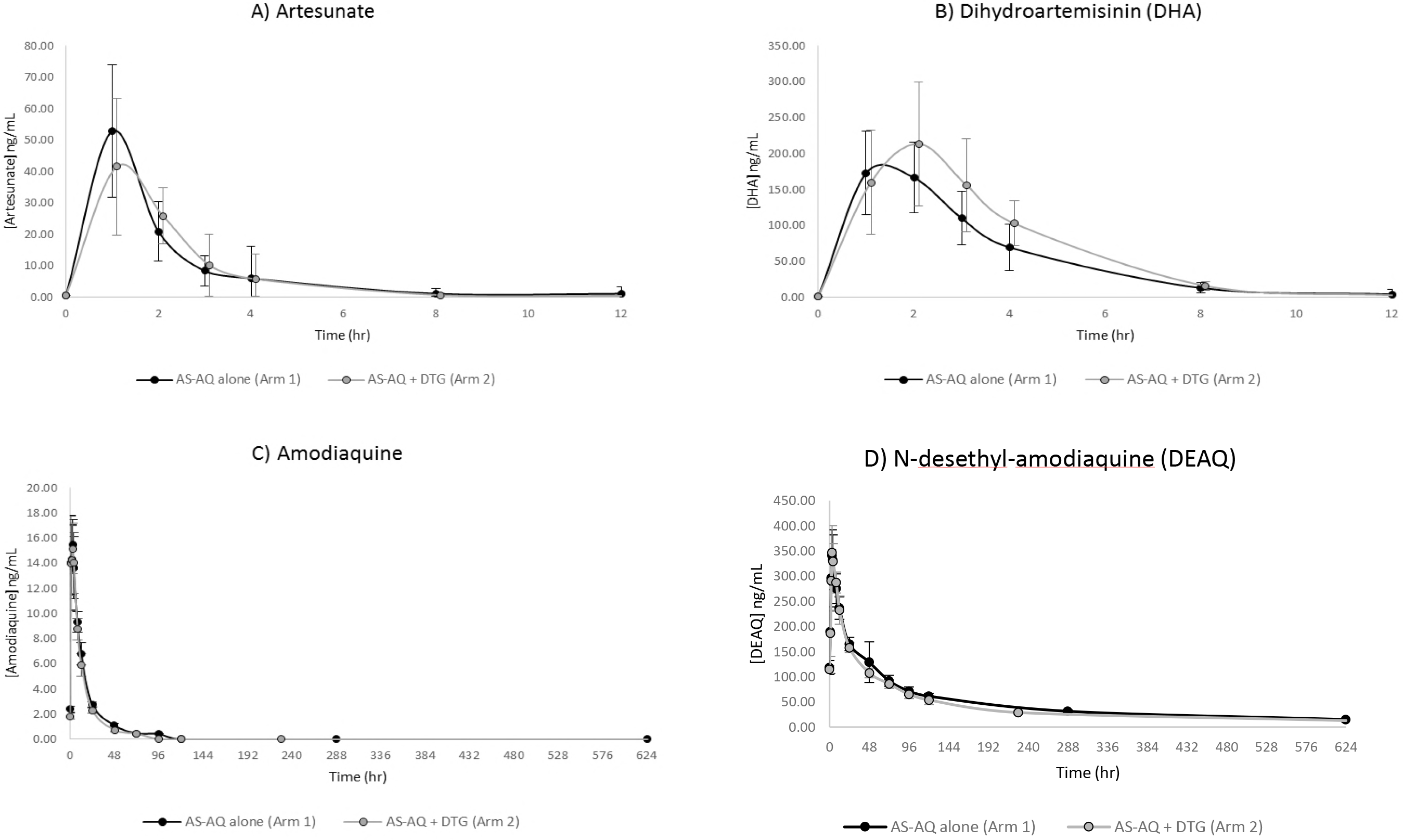
Artesunate-amodiauine parent and active metabolite PK.

